# Early-life environmental conditions shape demographic responses to climate change in male Antarctic fur seals

**DOI:** 10.64898/2026.06.01.729279

**Authors:** Anneke J. Paijmans, Jaume Forcada, Joseph I. Hoffman

## Abstract

Climate change intensifies selection, driving population declines and eroding genetic diversity, yet the mechanisms linking environmental perturbations to long-term demographic and evolutionary outcomes remain elusive. In particular, deteriorating early-life environments can have lasting impacts on fitness and population dynamics, but detecting these carry-over effects requires multidecadal, individual-based datasets. Long-term monitoring of Antarctic fur seals at Bird Island, South Georgia, has revealed a decline in breeding female numbers associated with persistent positive phases of the Southern Annular Mode (SAM), when warmer sea surface temperatures reduce local krill abundance. However, male responses remain understudied as their large size and aggressiveness preclude conventional tagging. To investigate male population dynamics, we combined 27 years of field observations of nearly 2,000 individually paint-marked territorial males with genetic recapture analysis of over 11,000 tissue samples, testing for both direct effects of the SAM on male abundance and lagged effects operating through recruitment. Territorial male numbers declined by 61%, yet male abundance was decoupled from the contemporary SAM, reflecting a delayed recruitment strategy that allows resource accumulation over multiple years and buffers short-term environmental variation. By contrast, recruitment probability fell by 59% annually and was strongly linked to natal conditions. Males born during increasingly positive SAM phases were significantly less likely to recruit, whereas heavier and more heterozygous individuals showed higher recruitment success, indicating that adverse natal environments act as selective filters. These findings identify a pathway linking environmental variability to population decline through recruitment dynamics, highlighting the importance of accounting for early-life carry-over effects when predicting the resilience of long-lived species to rapid environmental change.

## Introduction

Global climate change is creating novel selection pressures across taxa [1,2] forcing species to adapt or risk extinction [3,4]. These pressures can trigger rapid population declines, leading to demographic bottlenecks that erode genetic diversity through increased genetic drift and inbreeding [5–8]. Furthermore, deteriorating environmental conditions can act as a genetic filter, intensifying viability selection against individuals with genotypes of lower fitness and selectively removing them from the population, thereby compounding demographic and genetic erosion [9–11]. Nevertheless, the mechanistic pathways linking environmental change, altered selection pressures and population dynamics remain poorly understood, hindering our ability to predict biodiversity loss.

In particular, early-life environmental conditions can have lasting effects on life-history traits, shaping development, survival and reproductive strategies [12,13]. These effects may be immediate, such as when environmental change leads to increased neonatal mortality, or delayed, manifesting as a reduction in the probability of recruitment or diminished reproductive success later in life [14,15]. Importantly, when environmental conditions vary among cohorts, early-life effects can generate cohort-specific differences in individual quality that can influence population dynamics independently of the current environmental conditions [16]. However, establishing how anthropogenically altered early environments translate into long-term demographic outcomes in the wild is challenging because effects may only emerge years later, requiring multidecadal, individual-based studies that track environmental change and fitness across generations [17].

The Southern Ocean is one of the most rapidly changing marine environments on Earth, with rising sea surface temperatures, accelerating sea ice loss and shifts in primary productivity reshaping the entire ecosystem [18–20]. A key driver of this regional climatic variability is the Southern Annular Mode (SAM), the main mode of atmospheric circulation variability in the Southern Hemisphere. Over the past four decades, the SAM has shifted toward a positive phase, with recent values reaching levels unprecedented in the historical record [21]. Extended periods of positive SAM have reduced sea ice extent in the Antarctic Peninsula sector, negatively affecting the recruitment of Antarctic krill (*Euphausia superba*) and causing a poleward shift of the species’ range [18,22]. The resulting declines in local krill abundance have had cascading consequences for krill-dependent predators, contributing to reduced survival, breeding success and population growth across multiple seabird and pinniped species [23–27].

The Antarctic fur seal (*Arctocephalus gazella*) is a dominant Antarctic apex predator that has been strongly impacted by local declines in krill. Long-term monitoring of a study population at Bird Island, South Georgia, has shown that changes in the SAM and krill availability have intensified viability selection against homozygous female pups, resulting in a 17% increase in the mean heterozygosity of the breeding female population over two decades [10]. Concurrently, the female breeding population collapsed in 2009 and has continued to decline by around 7% annually since then [26], indicating strong demographic sensitivity to environmental change. Consistent with this, the number of breeding females arriving ashore each year closely tracks the contemporary environmental conditions, as only individuals with sufficient energetic reserves are able to successfully produce and rear a pup [10]. However, it remains unknown whether male Antarctic fur seals are affected by environmental change in the same way.

Juvenile Antarctic fur seal males are larger than females [28] and exhibit different foraging patterns [29,30], suggesting males may have an advantage in tracking the southward distribution of krill, potentially making them less susceptible to altered selection pressures under climate change. However, studying male Antarctic fur seals is challenging as their large size and aggressive behaviour preclude the use of conventional flipper tags and necessitate a combination of paint marking, remote biopsy sampling and genetic recapture analysis (i.e. the use of permanent genetic ‘marks’) to identify unique individuals both within and across years [31,32]. Furthermore, while females typically recruit into the breeding population at around 4–5 years of age, limited data from tagged male pups suggest that recruitment occurs much later at around 8–10 years of age [31]. Comprehensively documenting full male life-histories therefore entails decades of intensive fieldwork combined with the collection and analysis of individual-based observational and genetic data.

Here, we address this knowledge gap by investigating male Antarctic fur seal responses to climate change using 27 consecutive years of field observations of individual paint-marked territorial males together with genetic data from 11,276 tissue samples. Drawing on sightings of 1,955 unique adult males and genetic recapture analysis of 8,580 pups, we characterise changes in territorial male abundance over time and examine patterns of recruitment along with their potential drivers, including early-life environmental variation, birth mass and heterozygosity. By focusing on males, an understudied demographic in this system, we provide new insights into how early life carry-over effects shape the population dynamics of an important Southern Ocean predator in a rapidly changing environment.

## Results

We analysed 27 consecutive years of individual-based observational and genetic data from an intensively studied colony of Antarctic fur seals at Bird Island, South Georgia (Fig 1a). A total of 2,696 samples from adult males and 8,580 pups were genotyped at up to 39 microsatellite loci. These loci have previously been shown to be in Hardy-Weinberg equilibrium and linkage equilibrium in the study population and to capture variation in inbreeding [33–37].

**Fig 1.**
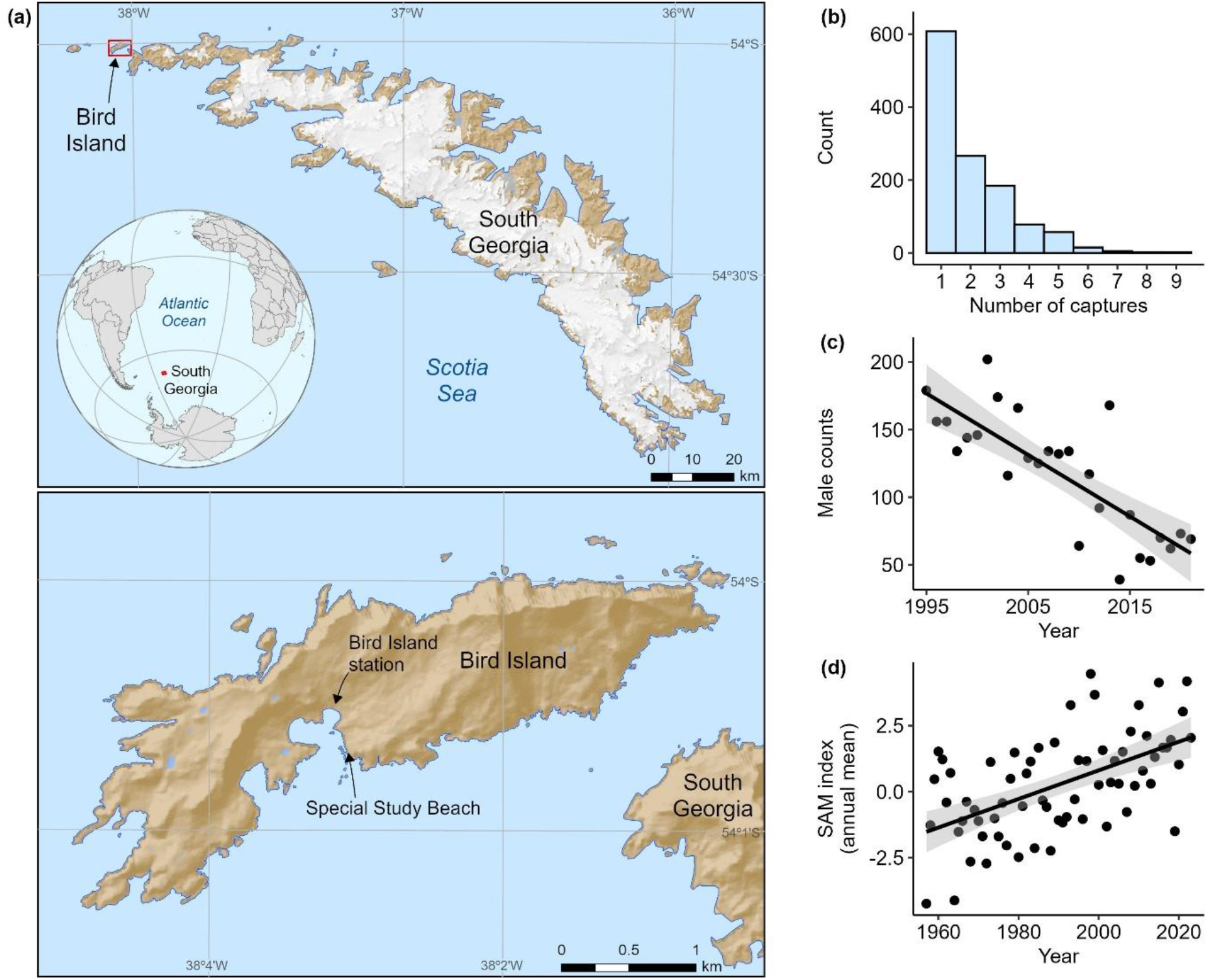
Study location and temporal patterns of seal abundance and environmental variation. (**a**) Map of South Georgia showing the location of the long-term study colony at Bird Island. Data from the South Georgia GIS (accessed 2025), produced by the Mapping and Geographic Information Centre, British Antarctic Survey, 2025; (**b**) The distribution of the number of years each territorial male was captured; (**c**) The temporal trend of territorial male counts; (**d**) The temporal trend of increasing annual mean Southern Annular Mode (SAM). The lines represent linear regressions fitted to the raw data (see Tables 1a and b for the parameter estimates) and the grey areas represent the associated 95% confidence intervals.

### Territorial male population trend

Using paint mark and genetic recapture data, we identified a total of 1,955 unique adult males that each held territories for between one and nine seasons (mean = two seasons, Fig 1b). The number of territorial males declined significantly over the past 27 years (GLM: *p* < 0.001; Fig 1c; Table 1a). Although the SAM increased significantly over the same period (LM: estimate = 0.054, s.e. = 0.010, t = 5.367, *p* < 0.001; Fig 1d) interannual variation in the SAM was not associated with the number of territorial males present in the study colony (GLM: *p* = ns; Table 1b).

**Table 1.**
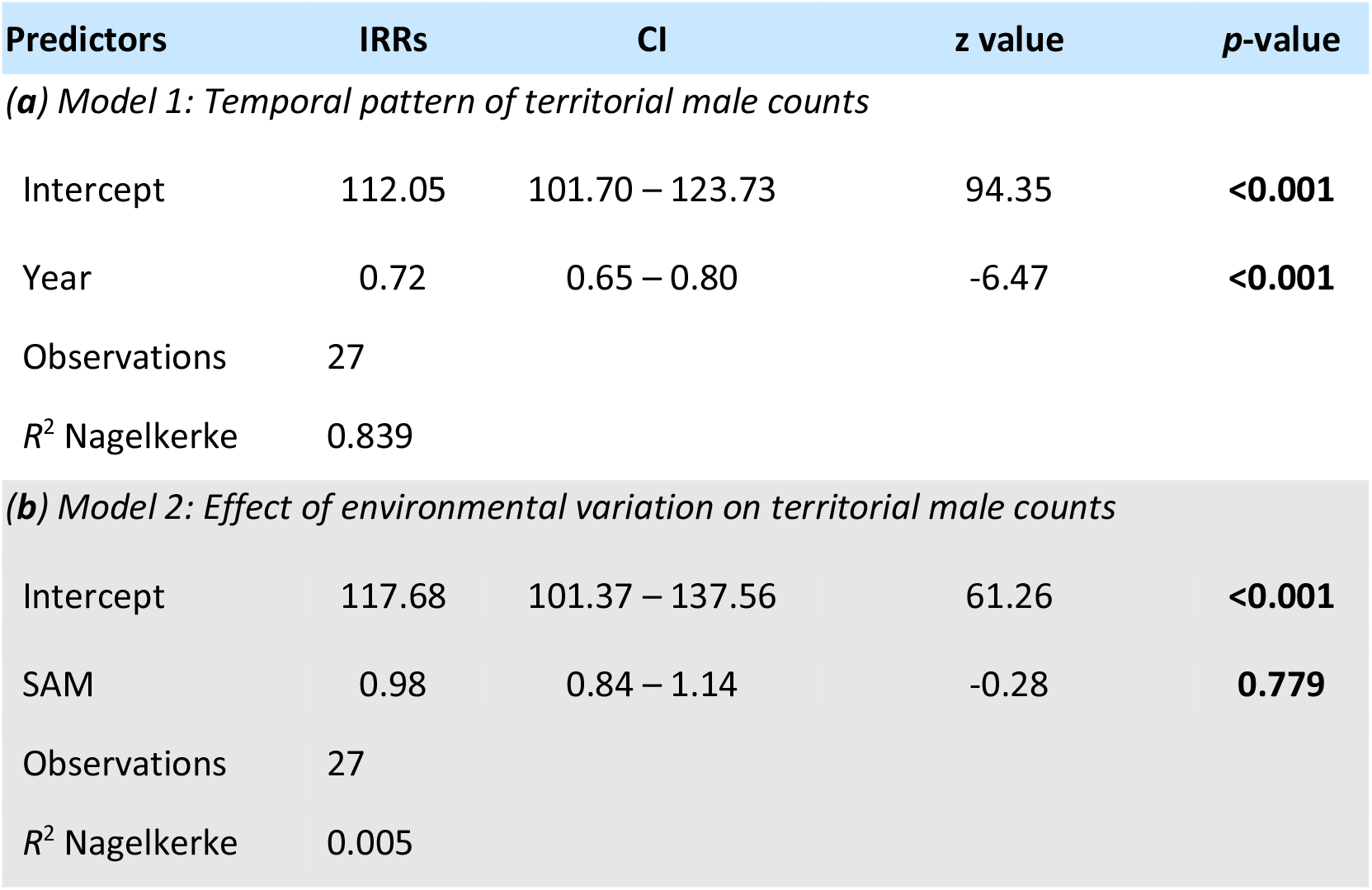
Parameter estimates from generalized linear mixed models with a negative binomial error distribution testing for the effects of (**a**) year and (**b**) annual mean SAM index on territorial male counts (see Materials and methods for details). Incidence rate ratios (IRRs) are shown together with 95% confidence intervals (CI). Significant *p*-values are highlighted in bold. The total number of observations and the variance explained (*R*^2^ Nagelkerke) are reported. Year and SAM were mean centered and scaled.

| Predictors | IRRs | CI | z value | *p*-value |
| --- | --- | --- | --- | --- |
| *(a) Model 1: Temporal pattern of territorial male counts* |  |  |  |  |
| Intercept | 112.05 | 101.70 – 123.73 | 94.35 | **<0.001** |
| Year | 0.72 | 0.65 – 0.80 | -6.47 | **<0.001** |
| Observations | 27 |  |  |  |
| $R^2$ Nagelkerke | 0.839 | | | |
| *(b) Model 2: Effect of environmental variation on territorial male counts* |  |  |  |  |
| Intercept | 117.68 | 101.37 – 137.56 | 61.26 | **<0.001** |
| SAM | 0.98 | 0.84 – 1.14 | -0.28 | **0.779** |
| Observations | 27 |  |  |  |
| $R^2$ Nagelkerke | 0.005 | | | |

### Patterns of male recruitment

Genetic recapture analysis of 8,580 pups identified 106 individuals that recruited into the territorial male population (Fig 2a). Consistent with the low *P*_ID_ of the genetic marker panel, which minimises the risk of spurious genetic matches, all of the recruited pups were male. No genetic matches were found between female pups and adult males, confirming the accuracy of the genetic recapture analysis and indicating that spurious matches are unlikely. The average age at recruitment was nine years (Fig 2b), with no significant difference among cohorts in recruitment age before versus after the 2009 population collapse (LM: estimate = 0.099, s.e. = 0.120, t = 0.828, *p* = ns). Consequently, pups born during the final nine years of the study (2013–2021 inclusive) were excluded from further analysis. Although recruits from these cohorts could be identified if they entered the territorial male population during the study period, the status of individuals not observed recruiting remains uncertain as some might still recruit after the study ended. Excluding these cohorts is therefore a conservative measure because it avoids prematurely classifying individuals that may still recruit after the end of the study as non-recruits. Taking into account all genotyped male pups up until 2013, we identified 106 recruited male pups and 2,273 non-recruited pups, corresponding to a recruitment rate of 4.5%.

**Fig 2.**
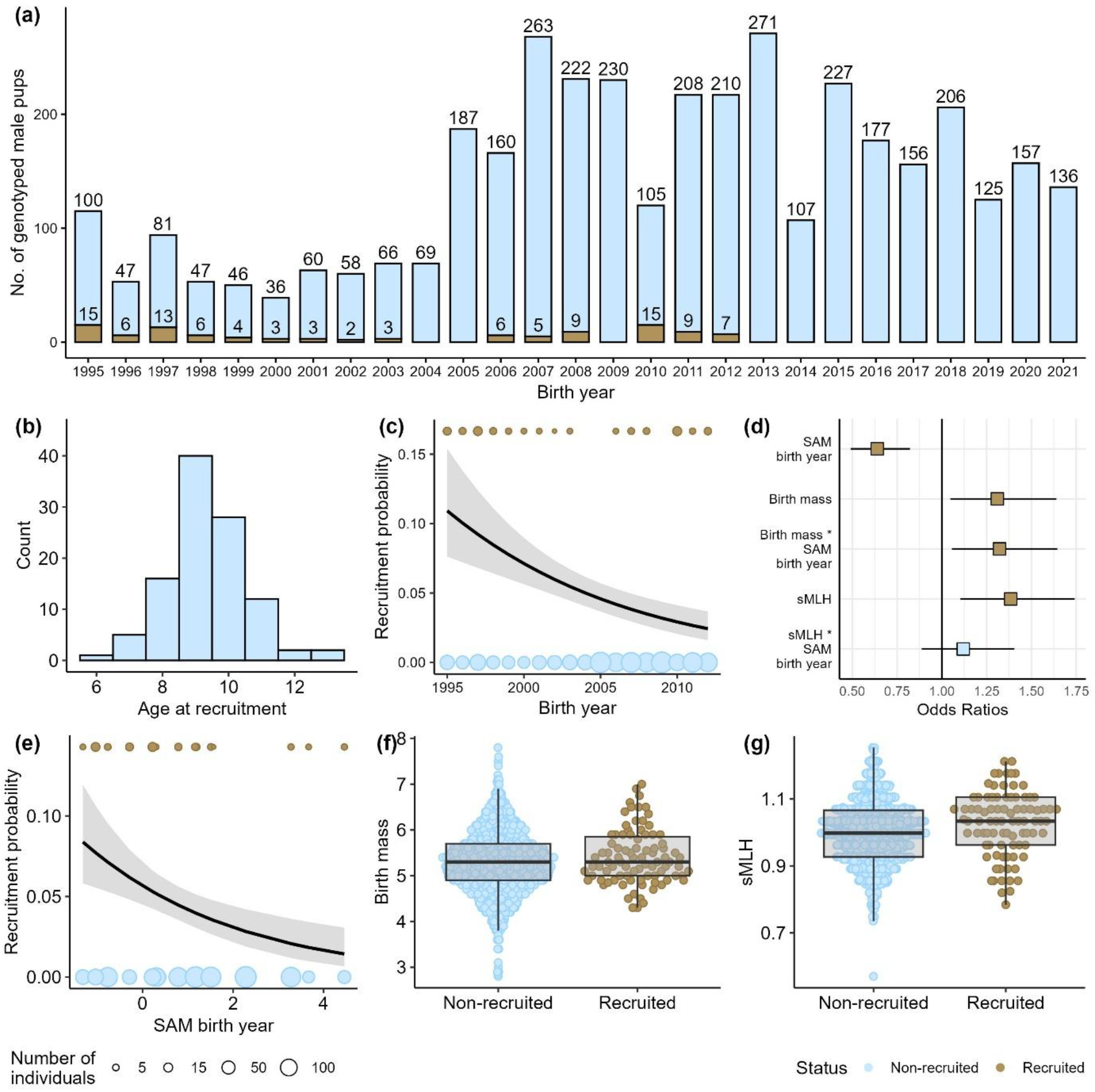
Long-term male recruitment dynamics. (**a**) Number of male recruits (brown) and non-recruits (blue) genotyped at 39 microsatellite loci per breeding season (total *n* = 3,863); (**b**) Distribution of age at recruitment for 106 genetically recaptured male pups; (**c**) temporal trend of recruitment probability. The model prediction is shown by the black line and the 95% confidence interval is shown by the grey shaded area. Point sizes scale in proportion to the number of individuals (brown = recruits and blue = non-recruits); (**d**) Model estimates (odds ratios) and associated 95% confidence intervals for the fixed effects of environmental, phenotypic and genetic predictor variables on male recruitment success. Statistically significant relationships are shown in brown (the parameter estimates are given in Tables 2a and b); (**e**) model prediction (black line) and 95% confidence interval (grey shaded area) for the relationship between recruitment probability and the Southern Annular Mode (SAM) of the birth year. Point sizes scale in proportion to the number of individuals (brown = recruits and blue = non-recruits); (**f**) Differences in birth mass between recruits (brown) and non-recruits (blue); (**g**) Differences in standardised multilocus heterozygosity (sMLH) between recruits (brown) and non-recruits (blue). The raw data are shown together with boxplots (centre line = median, bounds of box = 25th and 75th percentiles, upper and lower whiskers = largest and smallest value but no further than 1.5 * inter-quartile range from the hinge).

In parallel with the temporal decline in territorial male counts, we observed a significant decrease in male recruitment success over the past 27 years (GLM: *p* < 0.001; Table 2a), with the model predicting a 59% decrease in recruitment probability per year (Fig 2c). To investigate the genetic, phenotypic and environmental factors influencing recruitment, we constructed a second binomial GLM, again coding recruits as 1 and non-recruits as 0. Microsatellite heterozygosity (sMLH) was included as a predictor variable due to its known influence on female recruitment [10] together with birth mass, which affects survival in this and other pinniped species [34,38–40]. To evaluate the impact of early-life environmental conditions, we also fitted the SAM of the birth year as a predictor alongside two interaction terms: (i) SAM x sMLH, to test for environment-dependent inbreeding depression, and (ii) SAM x birth mass, to evaluate whether the benefits of higher birth mass depended on the environmental conditions at birth. Because our modelling framework excludes observations with missing data, these analyses was based on a reduced dataset of 87 recruits and 1,680 non-recruits with complete data.

**Table 2.** Parameter estimates from the generalized linear mixed models with a binomial error distribution testing for the effect of (**a**) year and (**b**) SAM index (annual mean), birth mass and heterozygosity on male recruitment success. Odd ratios are shown together with 95% confidence intervals (CI). Significant *p*-values are in bold. Total number of observations, as well as the variance explained (*R*^2^ Tjur) are reported. All predictors were mean centered and scaled.

| Predictors | Odds Ratios | CI | z value | p-value |
| --- | --- | --- | --- | --- |
| *(a) Model 1: Temporal effect on male recruitment success* |  |  |  |  |
| Intercept | 0.05 | 0.04 – 0.06 | -25.51 | **<0.001** |
| Year | 0.63 | 0.51 – 0.77 | -4.48 | **<0.001** |
| Observations | 1767 |  |  |  |
| $R^2$ Tjur | 0.014 | | | |
| *(b) Model 2: Environmental and genetic effect on male recruitment success* |  |  |  |  |
| Intercept | 0.04 | 0.03 – 0.06 | -24.15 | **<0.001** |
| SAM birth year | 0.64 | 0.49 – 0.82 | -3.41 | **0.001** |
| Birth mass | 1.31 | 1.05 – 1.64 | 2.36 | **0.018** |
| Birth mass * SAM birth year | 1.32 | 1.06 – 1.64 | 2.47 | **0.014** |
| sMLH | 1.38 | 1.10 – 1.74 | 2.79 | **0.005** |
| sMLH * SAM birth year | 1.12 | 0.89 – 1.40 | 0.96 | 0.337 |
| Observations | 1767 |  |  |  |
| $R^2$ Tjur | 0.016 | | | |

We found a significant negative effect of the SAM of the birth year on male recruitment success (GLM: *p* < 0.01; Fig 2d; Table 2b), with the model predicting a 56% decrease in recruitment probability with each unit increase in the SAM (Fig 2e). The most likely explanation for this pattern is that male pups born under adverse environmental conditions fail to survive beyond weaning, leaving no opportunity for recruitment. Accordingly, birth mass also had a positive effect on male recruitment success (GLM: *p* < 0.05; Fig 2d, f; Table 2b) and the interaction between birth mass and SAM was also significant (GLM: *p* < 0.05; Fig 2d; Table 2b), indicating that heavier pups have a greater advantage in poor years. Heterozygosity likewise had a significant positive effect on recruitment (GLM: *p* < 0.01 Fig 2d, g; Table 2b), but its interaction with SAM was not significant (GLM: *p* = ns; Fig 2d; Table 2b), suggesting that the benefits of high heterozygosity may not be environment-dependent, or alternatively that limited statistical power prevented us from detecting an interaction.

## Discussion

Understanding how environmental change translates into long-term demographic declines remains a central challenge in ecology and conservation. Although climate-driven shifts in marine ecosystems are well documented, direct evidence linking early-life conditions to recruitment and population dynamics is scarce due to the substantial logistical effort required to assemble individual-based datasets spanning multiple decades. To bridge this gap, we analysed 27 years of longitudinal and genetic data from a breeding colony of Antarctic fur seals at South Georgia. We detected a 61% decline in territorial male numbers, a trend attributable to a reduction in recruitment success linked to increasingly adverse early-life environmental conditions. Our findings demonstrate that changing shifting selection pressures during early life can have lasting demographic consequences for wild populations.

### Male population trend

Previous studies have shown that reproduction in female Antarctic fur seals is strongly tied to the current environmental conditions, resulting in a pronounced negative relationship between the number of females that come ashore to breed in a given year and the SAM [10]. By contrast, males show no such relationship, reflecting the distinct life-history strategies of the two sexes. Male Antarctic fur seals delay recruitment, allowing them to accumulate resources over multiple years to compete for territories. This delayed strategy appears to buffer them against short-term environmental fluctuations, decoupling their reproductive activity from the immediate effects of the SAM. Female Antarctic fur seals, by contrast, have relatively long reproductive lifespans of up to 18 years, promoting a trade-off between current and future reproductive investment and rendering them particularly sensitive to the immediate environmental conditions. In years of low food availability, many females demonstrably reduce reproductive investment or skip breeding altogether, presumably to conserve energy for future seasons. Males, by contrast, typically hold territories for only one or a few seasons but can sire large numbers of offspring [31]. Consequently, whereas female reproduction is constrained by current food availability, males rely on energy reserves accumulated over many years to achieve high reproductive success even under unfavourable environmental conditions.

### Male recruitment success

The male recruitment rate, estimated from genetic recapture analysis revealing 106 recruited and 2,273 unrecruited pups, was approximately 4.5%, roughly one third of the equivalent female recruitment rate of 16.3% (J. Forcada pers. comm.). This difference between the sexes may partly reflect the species’ polygynous mating system, in which only a small proportion of males successfully establish and maintain territories [31]. Under such intense male-male competition, some males might survive to adulthood but fail to recruit into the territorial male population because they are unable to secure or defend a territory. Alternatively, male pinnipeds are generally less philopatric than females [41,42], raising the possibility that some males may recruit at other colonies. However, individuals from the study colony are rarely observed breeding elsewhere around Bird Island and gene flow is restricted across larger spatial scales [33,43,44]. Dispersal is therefore unlikely to account for the low male recruitment rate, although the possibility of movements to unsampled sites around the South Georgia coastline cannot be excluded.

Importantly, male recruitment in our study is not measured conditional on survival. Direct data on juvenile survival in male Antarctic fur seals are lacking, but two lines of evidence suggest that early-life survival is likely to be a major factor limiting recruitment. First, apparent first year survival of female pups at the study colony has declined sharply over the past decade [26], a trend mirrored at distant sites including the South Shetlands [45]. This concordant pattern indicates that male pups may have experienced similar reductions in early-life survival. Second, among males born in years of high SAM, heavier pups were more likely to recruit. This is consistent with a general pattern among pinnipeds in which heavier pups have higher juvenile survival probabilities [10,34,46], again implying that viability selection during early life may be an important determinant of recruitment. By implication, most non-recruits likely died before reaching breeding age rather than surviving but failing to establish territories.

Further analysis revealed that the SAM of the birth year is a key environmental determinant of male recruitment success, with males born in low SAM years having a higher likelihood of recruiting into the breeding population. This pattern again suggests a mechanistic link to early survival: pups born under favourable environmental conditions may benefit from higher maternal investment during lactation and greater access to food following nutritional independence, both of which can increase the probability of surviving to adulthood [47,48]. More broadly, our results also align with the concept of “silver spoon effects”, where early-life advantages such as high quality environmental conditions at birth or greater maternal investment confer lasting fitness benefits [12–15,49]. These advantages could potentially extend beyond survival to influence the expression of traits such as body size and territorial behaviour later in life, with possible downstream effects on male reproductive success and the intensity of sexual selection.

Finally, theory predicts that inbreeding depression should be strongest early in life, as deleterious alleles are expressed and subsequently purged during early developmental stages, reducing their effects later in life [50,51]. However, several previous studies of Antarctic fur seals found no evidence for heterozygosity-fitness correlations in pups [34,35,52], suggesting that mortality at this stage is mainly caused by non-genetic factors such as trampling and starvation. This may either obscure underlying genetic effects or reflect weak selection on genetic quality during this life-history stage, when survival is strongly influenced by maternal provisioning and stochastic factors.

In this context, our finding that recruited male pups were significantly more heterozygous than non-recruits suggests that inbreeding depression is more strongly expressed during the transition from nutritional independence to recruitment, when individuals experience viability selection outside of the colony environment. A comparable pattern has been reported in female Antarctic fur seals, where deteriorating SAM conditions are associated with stronger viability selection against homozygous pups prior to recruitment [10]. Although we did not detect a significant interaction between heterozygosity and SAM of the birth year in males, this likely reflects a lack of statistical power due to low male recruitment rather than a genuine absence of environment-dependent inbreeding depression. Ultimately, consistent effects of heterozygosity on recruitment in both sexes align with the “early-life bottleneck hypothesis”, which argues that intense selection during early life ensures that the few individuals reaching the breeding population are those with the highest genetic resilience [50,51].

### Caveats

Because male Antarctic fur seals are too large and aggressive to be captured and tagged, we used a combination of paint marks and genetic recaptures to identify individuals both within and across years and to identify recruitment events. The probability of identity and the genotyping error rate for our dataset were both very low and no female pups were ever recaptured as adult males, suggesting that spurious matches were effectively absent. Nonetheless, we acknowledge several potential sources of bias. First, the loss of paint marks within seasons could inflate male counts if some animals are re-marked and recorded as new individuals. However, this effect is likely minimal as within-season recaptures are infrequent relative to between-season recaptures, indicating that paint marks are generally retained. Moreover, with approximately 80–90% of territorial males being sampled each season, the majority of paint mark loss events would be detected genetically. Second, because only sampled territorial males can be identified as recruits, any pups that recruited but were not sampled as adults would go undetected, potentially leading to slight underestimation of the recruitment rate. However, the high sampling coverage of territorial males suggests that this bias is unlikely to be of major importance. Third, genotyping errors could potentially split a single male into multiple apparent individuals, inflating the number of unique individuals and downwardly biasing our estimate of the recruitment rate. However, the genotyping error rate for our dataset was very low and all mismatching genotypes were carefully inspected to minimise this source of error in the genetic recapture analysis. In addition, the use of consistent field and laboratory protocols over several decades gives us no reason to believe that any of these biases would introduce systematic temporal trends into either male counts or recruitment estimates.

### Implications for population dynamics

Our study provides rare, multidecadal evidence linking climate change to demographic decline via early-life effects on male recruitment. When conditions experienced early in life have stronger effects on survival than conditions experienced later in life, population dynamics can be disproportionately influenced by lagged cohort effects [16]. In other words, the number of territorial males present in any given year is not solely determined by that year’s environment, but reflects the cumulative impact of previous years in which cohorts of males experienced adverse early-life conditions. Over time, this creates a delayed response in the territorial male population, where the effects of adverse environmental conditions during sensitive environmental phases propagate across multiple generations, amplifying the long-term decline in territorial male numbers.

Sex-specific differences may further complicate this picture. If males and females respond differently to early-life environmental conditions, population dynamics could become sex-biased. In polygynous systems, reduced male recruitment may have limited short-term effects on population growth, since relatively few males are required for successful reproduction. However, sustained reductions in male recruitment could, over time, alter the mating system. A previous study of a low-density Antarctic fur seal breeding colony at the South Shetland Islands reported even stronger male reproductive skew than at Bird Island, with the presence of multiple full siblings across years indicating that parents remated across consecutive breeding seasons [53]. This suggests that dominant males may monopolise access to sexually receptive females more effectively at low population density. Consequently, a decline in population density at South Georgia could potentially increase the strength of polygyny, reducing the effective population size and heightening the risk of inbreeding. Future work should investigate temporal changes in the mating system and their consequences for genetic diversity and evolutionary potential.

## Conclusions

This study shows that variation in early-life environmental conditions is a major driver of male recruitment in Antarctic fur seals and has contributed substantially to the long-term decline in the number of territorial males at our long-term monitoring site in South Georgia. By implication, environmental variability experienced during early life can have enduring demographic consequences, with population trajectories reflecting cumulative lagged impacts across multiple cohorts rather than responding immediately to short-term environmental variation. Understanding how early-life effects carry over to influence male reproductive performance will help to further unravel the mechanisms shaping population change in this iconic marine predator and inform evidence-based management of the Southern Ocean ecosystem.

## Methods

### Field methods

This study was conducted at an Antarctic fur seal breeding colony on Bird Island, South Georgia (Fig 1a, 54°00024.800 S, 38°03004.100 W) during the austral summers of 1994/1995 to 2020/2021 inclusive (breeding seasons are hereafter referred to by the year in which they ended). The study colony was separated from adjacent breeding sites by a cliff on the east side, open sea on the west, and rocky ridges to the north and south. A scaffold walkway [54] provided access to all parts of the colony while minimising disturbance to the animals and maximising the safety of the researchers.

The seals were captured and restrained following protocols from the British Antarctic Survey (BAS) long-term monitoring and survey program. Over the course of the study, over a thousand adult females were tagged using cattle ear tags (Dalton Supplies, Henley on Tames, UK) placed in the trailing edge of the forefipper. Pups born to these females were captured on the day of birth, their sex and mass were recorded, and a tissue sample was collected from the interdigital margin of the foreflipper using piglet ear notching pliers [55]. In addition, a variable number of pups with unknown mothers were sampled each year, depending on the available field staff resources, either at birth or towards the end of the field season when the pups were recaptured for permanent tagging.

Adult males could not be captured and tagged due to their large size and aggressiveness. Hence, males occupying territories within the colony were given individually distinctive paint marks for within-season identification and were tissue sampled using a remote biopsy dart system [56]. Daily surveys were then made of all territorial males from 1^st^ November until the end of the pupping period (early January). All tissue samples were stored individually at -20°C in 20% dimethyl sulphoxide (DMSO) saturated with salt [57].

### Microsatellite genotyping

Total genomic DNA was extracted using an adapted phenol-chloroform protocol [58] and all of the samples were genotyped at between nine and 39 microsatellite loci. Samples collected between 1994 and 2008 inclusive were genotyped at nine microsatellites as described by Hoffman and Amos [59]. Briefly, polymerase chain reactions (PCRs) were performed separately for each locus while incorporating [α^32^P]-dCT. The PCR products were then resolved by electrophoresis on standard 6% acrylamide sequencing gels and detected by autoradiography. The autoradiographs were manually scored and any reactions yielding uncertain genotypes (e.g. with faint or unclear banding patterns) were repeated.

Samples collected during and after 2009 were genotyped at 39 microsatellite loci as described by Paijmans et al. [33]. Briefly, the microsatellites were PCR amplified in five separate multiplex reactions using a Type It Kit (Qiagen). The resulting fluorescently labelled PCR products were resolved by electrophoresis on an ABI 3730xl capillary sequencer (Applied Biosystems, Waltham, MA, USA) and allele sizes were called relative to the LIZ size standard (1:150 dilution) using GeneMarker 2.6.2 (SoftGenetics, LLC., State College, PA, USA). To ensure high genotyping quality, all of the traces were manually inspected by at least two experienced observers, and any incorrect calls were corrected.

To achieve a seamless transition from manual to fluorescent genotyping, we carefully harmonised the allele scoring throughout the entire dataset. Internal consistency was checked by independently genotyping 48 randomly selected samples using both methods. For the nine shared loci, only 4 out of a total of 424 reactions yielded different genotypes, corresponding to an error rate of 0.0095 per reaction, all of which were due to scoring or data entry errors. Above and beyond this high level of internal consistency, genotyping error rates were consistently low for both methods, at 0.0013– 0.0074 per reaction for the manual approach [59] and 0.0050% per reaction for the fluorescent approach [33]. Our final dataset comprised a total of 8,580 unique pups and 2,696 adult male samples, of which 7,413 (86%) and 829 (31%) respectively were genotyped at 39 microsatellite loci.

### Genetic identity analyses

To identify adult males that were sampled more than once within or across years, we used a custom R script to find duplicate genotypes within our dataset of 2,696 adult male samples. Given that different individuals can have identical multilocus genotypes when too few loci are genotyped, we first calculated the probability of identity (*P*_ID_ [60]) across all individuals and loci. This was very low (9 loci = 1.29 × 10^−12^; 39 loci = 1.19 × 10^−43^) indicating that identical genotypes almost certainly represent resampled individuals. Because the dataset may include some related individuals, we also calculated the more conservative *P*_ID_ among siblings (*P*_ID Sib_ [61]). This was still low enough to be able to distinguish full siblings with high confidence (9 loci = 1.10 × 10^−4^; 39 loci = 5.44 × 10^−16^).

To account for genotyping errors, which can lead to overestimation of the number of individuals in a population [62,63], our custom script identified genetic matches while allowing a maximum of two mismatching loci out of nine (for comparisons where at least one sample was genotyped at nine loci) or seven mismatching loci out of 39 (for comparisons where both samples were genotyped at 39 loci). All of the mismatches were then visually inspected as described by Hoffman et al. [64]. Scoring and data entry errors were corrected, and when no mismatches remained, matching samples were assigned to the same individual. Samples with genuine mismatches were classified as belonging to different individuals. This yielded a total of 1,217 unique genotyped adult males that were recaptured between one and 14 times each (mean = two recaptures).

Finally, we sought to identify individuals born in the study colony that subsequently recruited into the adult territorial male population by comparing the genotypes of 8,580 pups (4,063 males, 4,040 females and 477 pups of unknown sex) with those of the 1,217 unique genotyped adult males. As described above, genetic matches were identified while allowing for mismatches, which were then visually inspected to identify genotyping errors and corrected where appropriate. This procedure identified a total of 106 recruited male pups.

### Data selection, additional genotyping and calculation of heterozygosity

To minimise missing data, we selected the genotype with the fewest missing loci for each recaptured individual to represent that individual. To maximise genetic resolution for estimating individual heterozygosity, all recruits and non-recruits sampled up to and including 2008, which were only genotyped at 9 loci, were retrospectively genotyped at 39 loci. Standardized multilocus heterozygosity (sMLH [65]) was then calculated for all individuals successfully genotyped for at least 28 loci (mean = 38.6 loci) using the *sMLH* function of the InbreedR package [66].

### Statistical analysis

To investigate temporal changes in the number of territorial males, we tallied the number of unique individuals observed ashore each season, integrating both paint mark and genetic. Duplicate samples of the same individuals within seasons were removed from the counts. We then constructed a generalised linear model (GLM) with a negative binomial error distribution, fitting male counts as the response variable and year as a continuous predictor variable. The model was specified as follows:

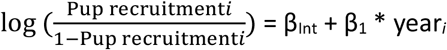

where:

*µ*_*i*_ represents the probability that recruitment is equal to 1 for the *i*-th individual in the sample, and β_Int_, β_1_ are regression coefficients for the intercept and the predictor variables, and ε is the random error.

To investigate the effect of environmental variation on male counts, we constructed a second GLM, fitting male counts as the response variable and the annual mean SAM index of the sampling year (available at https://legacy.bas.ac.uk/met/gjma/sam.html) as a continuous predictor variable. The model was specified as follows:

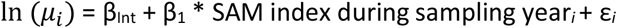

where:

*µ*_*i*_ represents the probability that recruitment is equal to 1 for the *i*-th individual in the sample, and β_Int_, β_1_ are regression coefficients for the intercept and the predictor variables, and ε is the random error.

To investigate temporal changes in male recruitment success, we constructed a GLM with a binomial error structure, coding the response variable as 1 for recruits and 0 for non-recruits. Because males recruit at an average age of nine years, we excluded the last nine seasons (2013–2021) from our model, as recruitment status could only be determined with certainty for cohorts that had sufficient time to recruit during the study period. This approach is conservative because it excludes pups that recruited after this cutoff. In addition, we assumed that age at recruitment remained constant across cohorts, while acknowledging that it may have shifted following the 2009 collapse; although, our results provide no evidence for this (see Results). Year was fitted as a continuous predictor variable. The model was specified as follows:

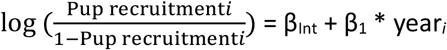

where: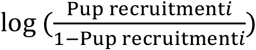 represents the probability that recruitment is equal to 1 for the *i*-th individual in the sample, and

β_Int_, β_1_ are regression coefficients for the intercept and the predictor variables.

To identify factors influencing male recruitment success, we constructed a second GLM with a binomial error structure, again coding the response variable as 1 for recruits and 0 for non-recruits and excluding the last nine seasons (2013–2021). We fitted sMLH and birth mass as continuous predictor variables. To test for the effects of early-life environmental conditions on recruitment success, we also fitted the SAM of the birth year and included two-way interactions between SAM and the other two predictors (sMLH and birth mass). The model was specified as follows:

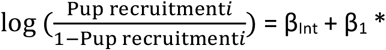 SAM index in birth year_*i*_ + β_2_ *birth mass_*i*_ + β_3_ *birth mass_*i*_ * SAM index in birth year_*i*_ + β_5_ * sMLH_*i*_ + β_5_ * sMLH_*i*_ * SAM index in birth year_*i*_

where:

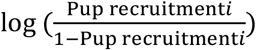 represents the probability that recruitment is equal to 1 for the *i*-th individual in the sample, and

β_Int_, β_1_–β_5_ are regression coefficients for the intercept and the predictor variables.

In all of the models, the predictors were centered by subtracting the means from the observed values and scaled by dividing the centered values by their standard deviations to improve model convergence. To verify that the model assumptions were met, we inspected the residuals for normality (Q-Q plots), linearity and equality of error variances (residuals versus predictions plots) and checked for under/overdispersion. These diagnostic checks were implemented using the DHARMa package [67].

## Ethics

Antarctic fur seal sampling and handling was carried out by the British Antarctic Survey and approved by the British Antarctic Survey Animal Welfare and Ethics Review Body (reference no. PEA6 and AWERB applications 2018/1050 and 2019/1058). Samples were collected as part of the Polar Science for Planet Earth programme of the British Antarctic Survey and under permits from the Government of South Georgia and the South Sandwich Islands (GSGSSI, Wildlife and Protected Areas Ordinance (2011), RAP permit numbers 2018/024 and 2019/032). The samples were imported into the UK under permits from the Department for Environment, Food and Rural Affairs (Animal Health Act, import license number ITIMP18.1397) and from the Convention on International Trade in Endangered Species of Wild Fauna and Flora (import numbers 578938/01-15 and 590196/01-18).

## Data accessibility

Microsatellite and fitness data can be accessed via Zenodo repository: https://zenodo.org/records/19680324 [68]. The SAM index data were downloaded from http://www.nerc-bas.ac.uk/icd/gjma/sam.html on the 3^rd^ of April 2024. All data wrangling steps and statistical analyses were implemented in R. The complete documented workflow for the genetic recapture analysis is available via our GitHub repository: https://github.com/apaijmans/AFS_recaptures/. The analyses of the temporal and environmental effects of male counts and recruitment are available via the GitHub repository: https://github.com/apaijmans/AFS_male_recruits/.

## Declaration of AI use

We have not used AI-assisted technologies in creating this article.

## Acknowledgements

We thank the British Antarctic Survey’s field assistants for the collection of tissue samples and observational data. We also thank Ira Wachendorf, Ane-Liv Berthelsen, Felicitas Christaller and Nicole Kröcker for help with genotyping and preliminary analyses.

## Authors’ contributions

A.J.P.: conceptualization, data curation, formal analysis, investigation, methodology, validation, visualization, writing—original draft, writing—review and editing;

J.F.: funding acquisition, investigation, methodology, project administration, resources, validation, writing—review and editing;

J.I.H.: conceptualization, data curation, formal analysis, funding acquisition, investigation, methodology, project administration, resources, supervision, validation, visualization, writing— original draft, writing—review and editing.

All authors gave final approval for publication and agreed to be held accountable for the work performed therein.

